## Supplementary Figures and Tables for "3D genomic analysis reveals novel enhancer-hijacking caused by complex structural alterations that drive oncogene overexpression"

Mortenson et al.

#### Supplementary Information

Included are 9 Supplementary Figures (including figure legends) and 3 Supplementary Tables.

### Supplementary Figure 1

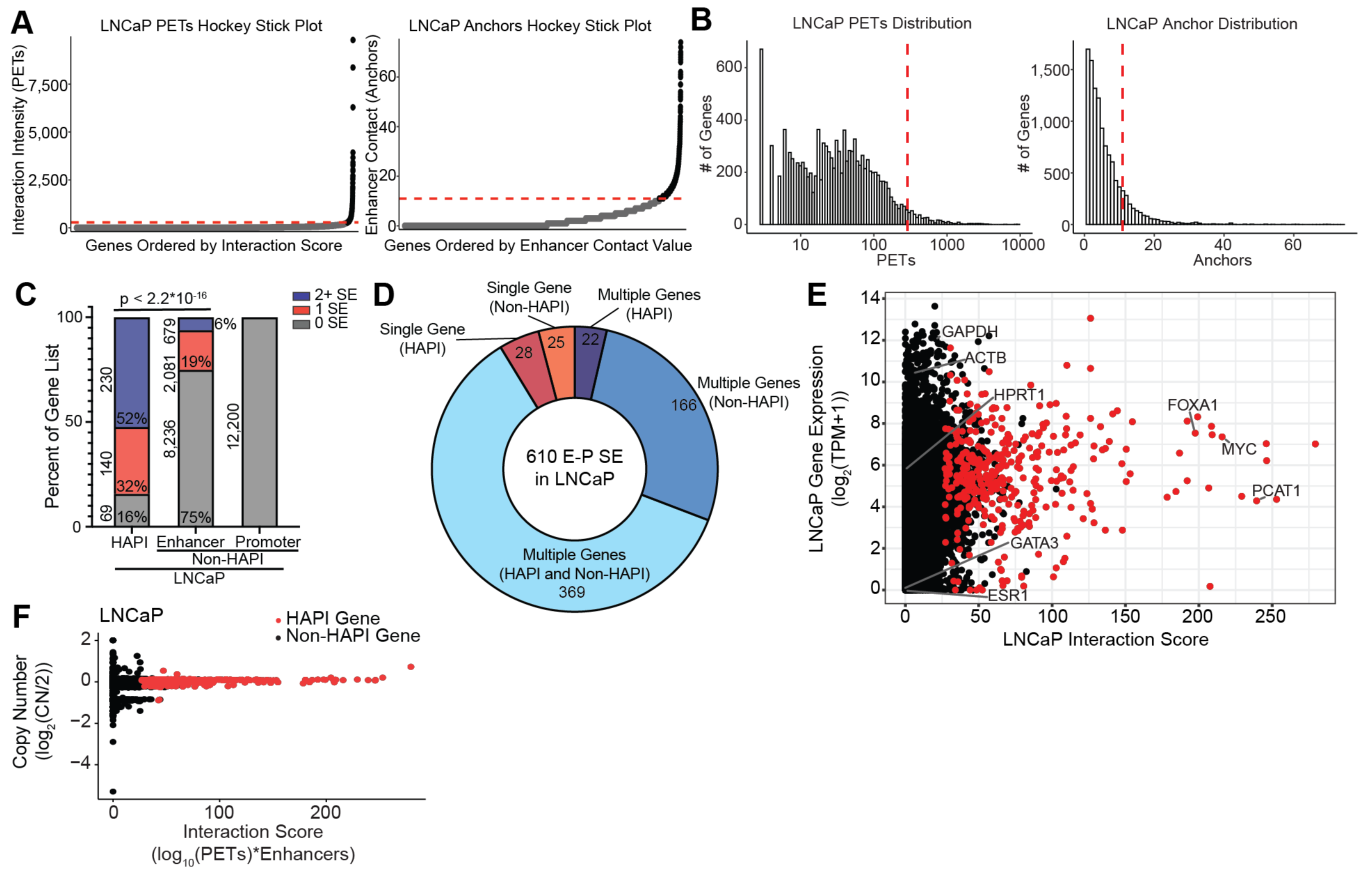


**Figure S1: Additional profiling of HAPI genes. A:** Hockey stick plots showing LNCaP interaction intensity and enhancer contact values of each gene in LNCaP cells, with inflection point cutoffs specified (red dashed lines). **B:** Distribution of interaction intensity and enhancer contact values for all genes in LNCaP with cutoffs from inflection points specified (red dashed lines). **C:** The percentage of HAPI genes and enhancer-connected or promoter-driven (no enhancer interaction) non-HAPI genes that are looped to super-enhancers. P-value was calculated using two-sided Fisher exact tests. **D:** The percentage of super-enhancers, involving enhancer-promoter (E-P) loops, that are looped to HAPI and/or non-HAPI genes in LNCaP cells. **E:** Interaction score versus log2(TPM+1) gene expression for HAPI genes (red) and non-HAPI genes (black) in LNCaP. **F:** Interaction score versus copy number (log2(CN/2)) for HAPI genes (red) and non-HAPI genes (black) in LNCaP.

#### Supplementary Figure 2

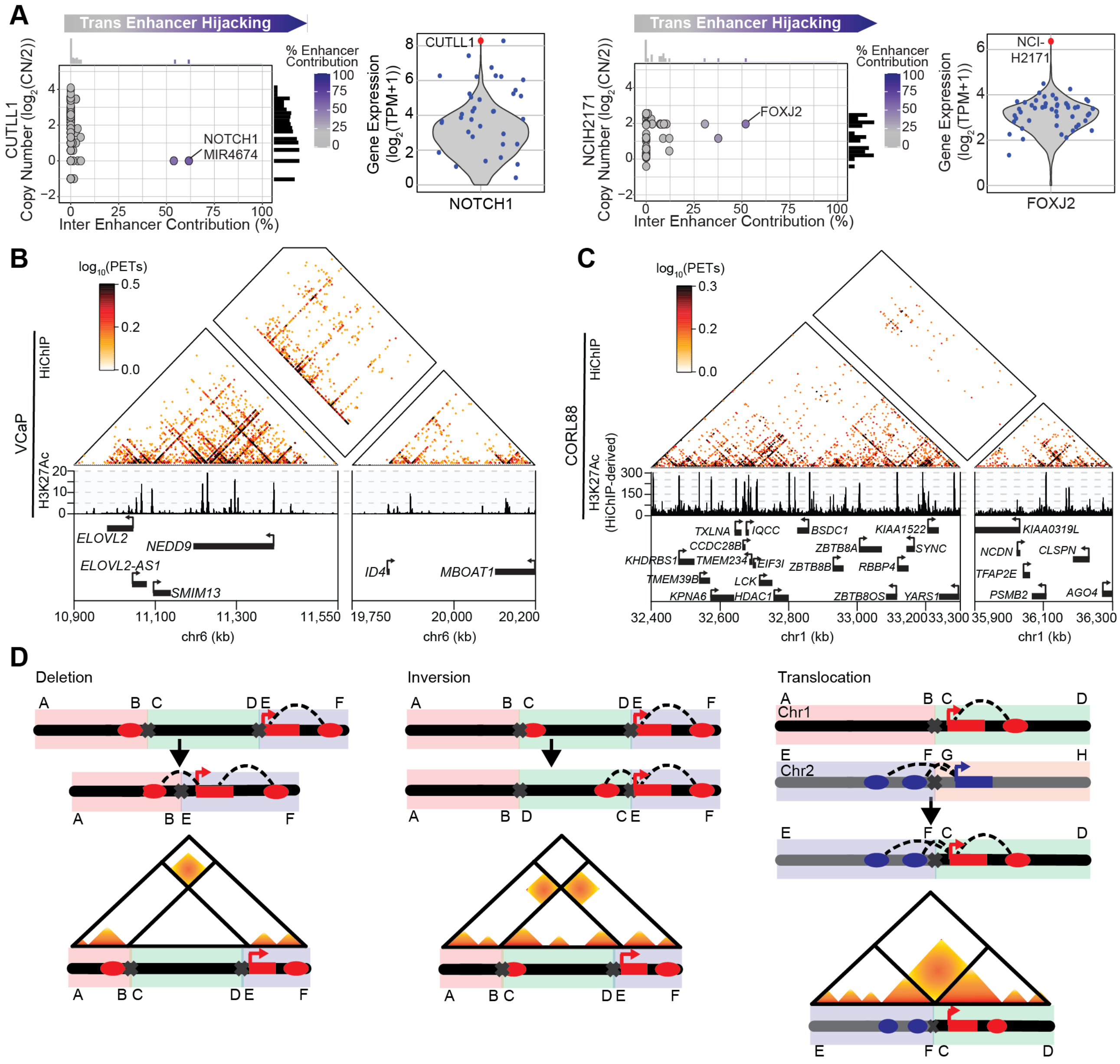


**Figure S2: Additional examples of enhancer-hijacking genes identified by HAPI analysis. A:** Inter enhancer contribution and copy number distribution of all HAPI genes for CUTLL1 and NCIH2171 with trans enhancer hijacking genes *NOTCH1* and *FOXJ2* (blue) labeled. Gene expression plots for the enhancer hijacking genes *NOTCH1* and *FOXJ2* generated using RNA-seq data with the identified cell lines CUTLL1 and NCIH2171 (red dots), respectively, compared to all cell lines of the same cancer type (blue dots) and all CCLE cell lines (grey violin plots). **B-C:** H3K27ac HiChIP and ChIP-seq signal at chromosomes 6 and 1 regions in VCaP (B) and CORL88 (C), respectively. The H3K27ac ChIP-seq signal in CORL88 is derived from HiChIP reads. **D:** Schematic showing different types of structural alterations, the resulting enhancer-promoter interactions, and how these structural variants would appear in HiChIP heatmaps.

#### Supplementary Figure 3

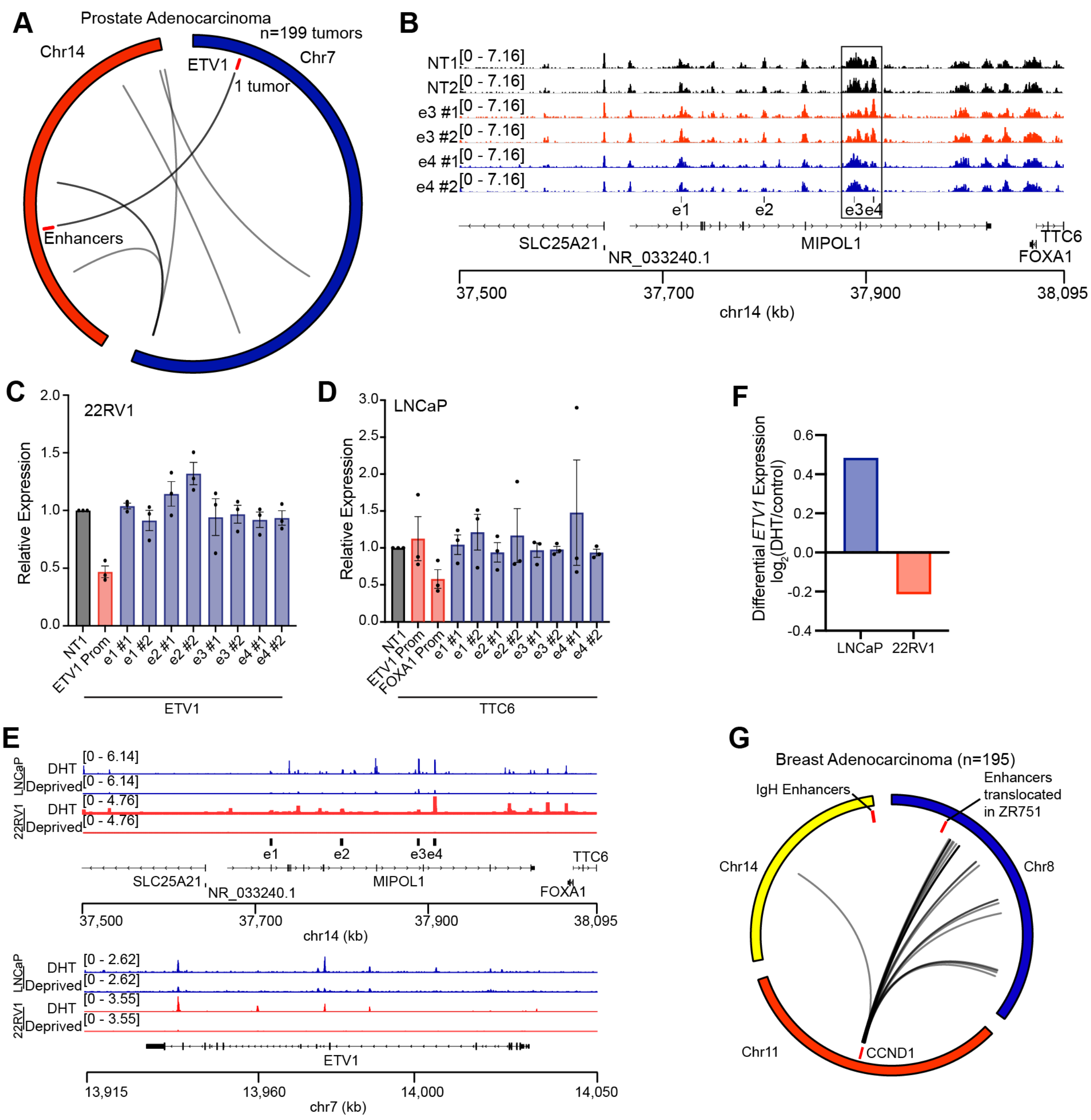


**Figure S3: Additional characterization of enhancer hijacking in LNCaP and tumor samples. A:** Circos plot displaying translocations between chromosomes 7 and 14 in 199 prostate adenocarcinoma tumor samples from PCAWG data set with *ETV1* and identified hijacked enhancers annotated. **B:** H3K27ac ChIP-seq in LNCaP-dCas9-KRAB-MeCP2 cells infected with non-targeting sgRNAs (NT1 and NT2) and enhancer targeting sgRNAs (e3 and e4). **C-D:** RT-qPCR measuring expression changes of *ETV1* in 22RV1 (C) and *TTC6* in LNCaP (D) after CRISPRi of each defined enhancer e1-e4 or the promoters of *ETV1* and *FOXA1/TTC6*. Gene expression levels were normalized to cells treated with the non-targeting sgRNA NT1. n = 3 biologically independent experiments. Data are presented as mean values +/- SEM. Source data are provided as a Source Data file. **E:** AR ChIP-seq in LNCaP and 22RV1 cells with and without DHT treatments for genomic regions containing the hijacked enhancers (top) and *ETV1* (bottom). **F:** log2-transformed fold changes of RNA-seq counts per million values for *ETV1* in LNCaP and 22RV1 cells with and without DHT treatments. **G:** Circos plot displaying translocations among chromosome 8, 11, and 14 (lines for translocations stemming from a 2 Mb window centered at *CCND1*) in 195 breast adenocarcinoma tumors samples from PCAWG data set with *CCND1* and identified hijacked enhancers annotated.

### Supplementary Figure 4

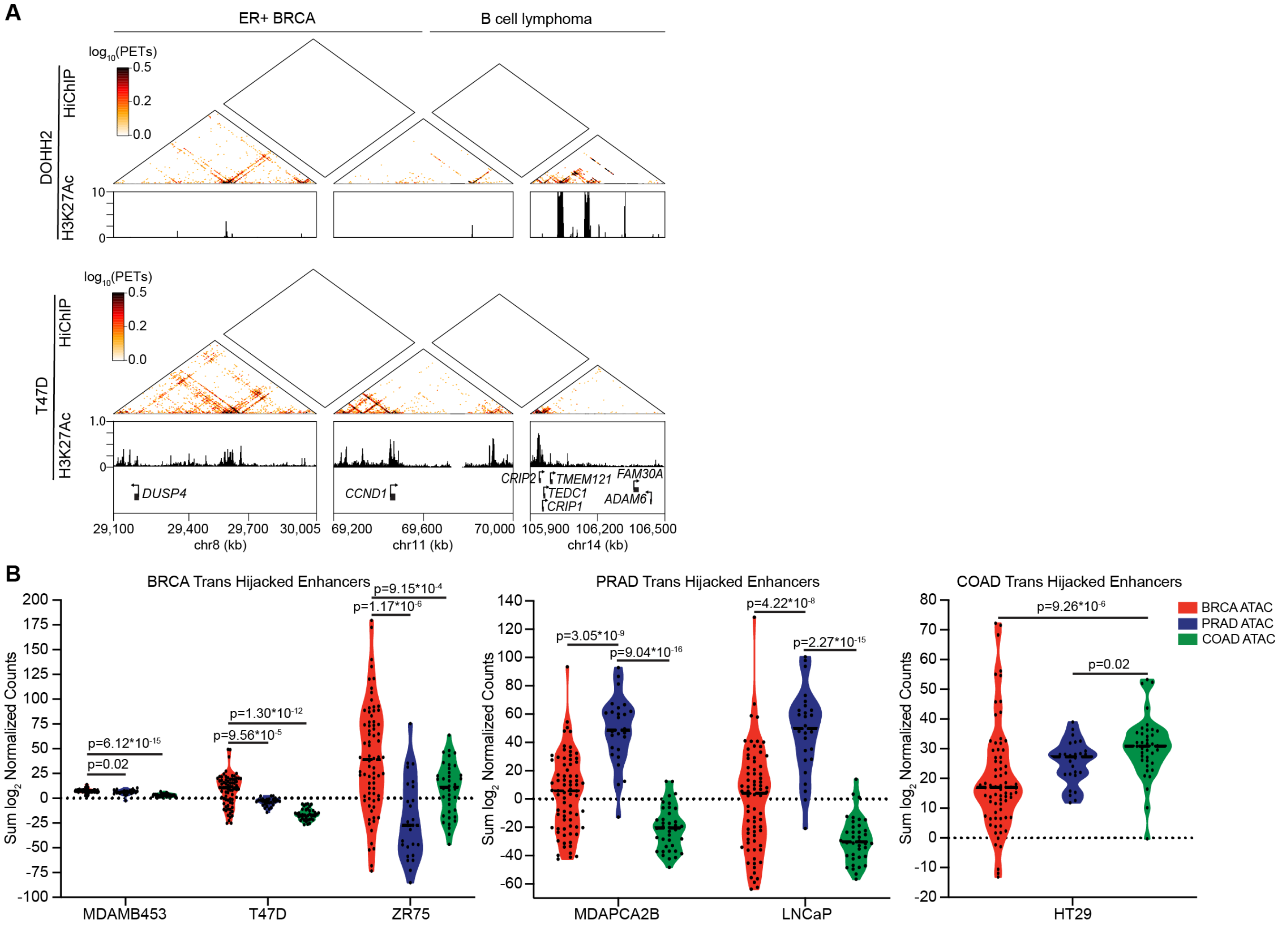


**Figure S4: Enhancer origin of hijacked enhancers is cancer-type specific. A:** DOHH2 and T47D H3K27ac HiChIP and ChIP-seq signal at the *CCND1* locus and the hijacked enhancer regions found in REC1 and ZR751 cells. **B:** Comparison of log2 normalized counts of the TCGA ATAC-seq peaks, overlapping with HAPI gene-hijacked enhancers in the indicated cell lines, for breast cancer (BRCA), prostate cancer (PRAD), and colorectal cancer (COAD) TCGA tumor tissues. P-values were calculated from two-sided wilcox tests.

### Supplementary Figure 5

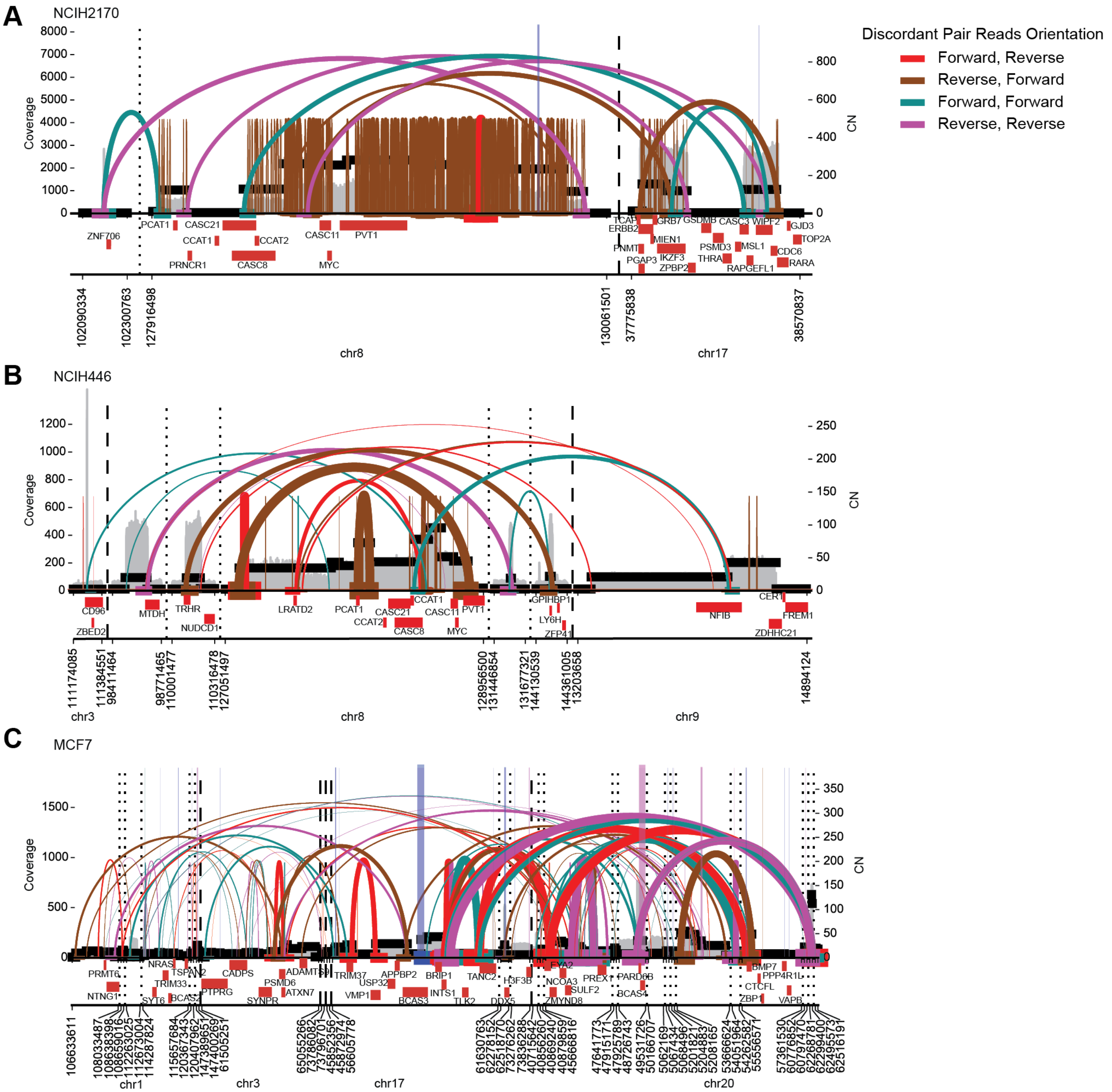


**Figure S5: AmpliconArchitect reconstructions for NCIH2170, NCIH446, and MCF7. A-C:** AmpliconArchitect plots depicting WGS-derived copy number and structural variants with labeled gene tracks for NCIH2170 (A), NCIH446 (B), and MCF7 (C).

#### Supplementary Figure 6

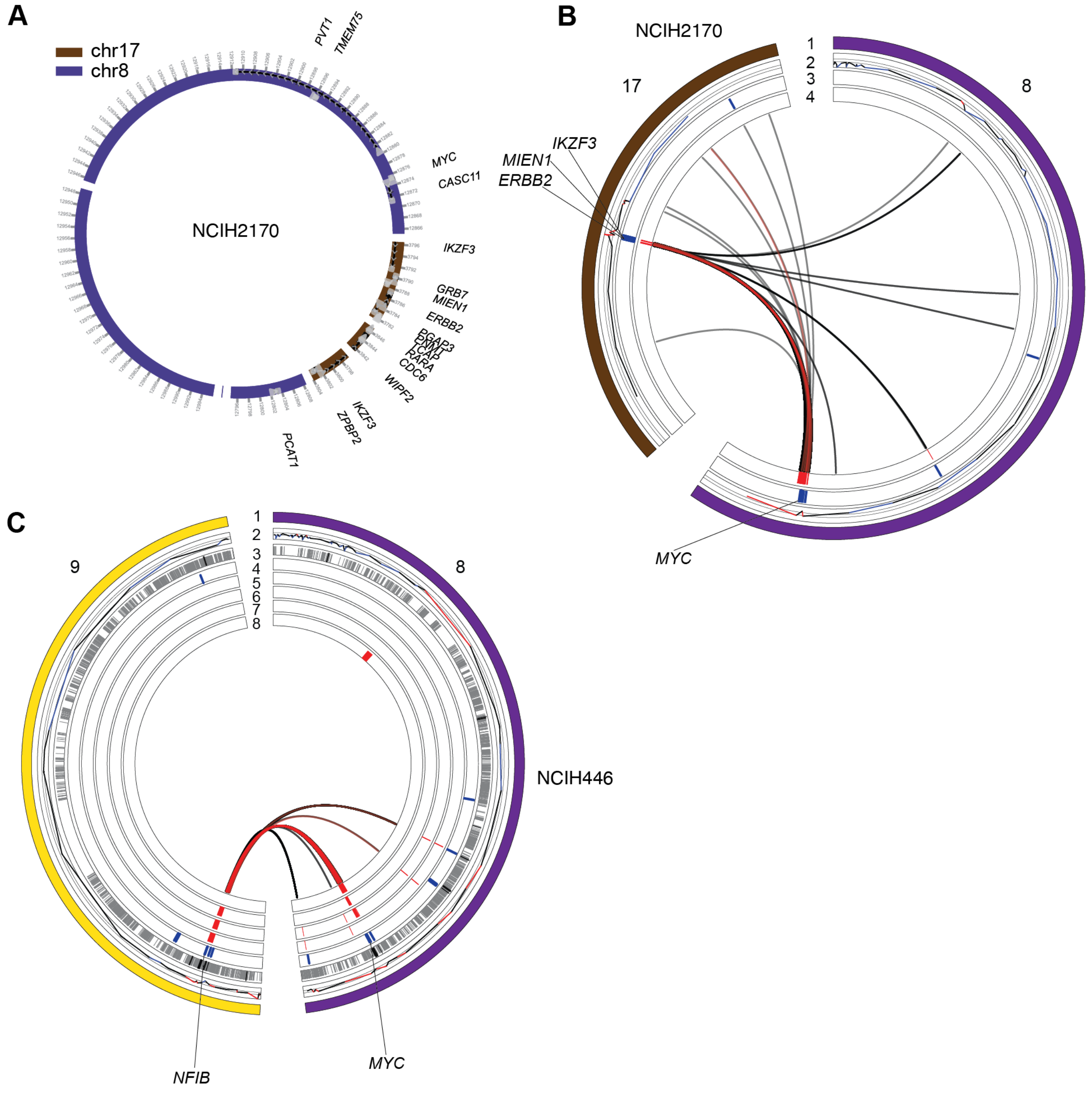


**Figure S6: Complex amplicons in NCIH2170 and NCIH446. A:** Structure of a proposed ecDNA in NCIH2170 based on AmpliconArchitect results. **B:** Circos plot of chromosomes 17 and 8 in NCIH2170 showing: 1) chromosome, 2) copy number, 3) TSS of HAPI genes in NCI2170, 4) DNA segments associated with the potential ecDNA, and in the center HiChIP enhancer-promoter interactions of HAPI genes (black), and WGS translocations (red). **C:** Circos plot of chromosomes 9 and 8 in NCIH446 showing: 1) chromosome, 2) copy number, 3) H3K27ac peaks showing enhancers (grey) and super-enhancers (black), 4) TSS of HAPI genes in NCIH446, 5-8) DNA segments of AmpliconArchitect-predicted amplicons, and in the center HiChIP enhancer-promoter interactions of HAPI genes (black), and WGS translocations (red).

#### Supplementary Figure 7

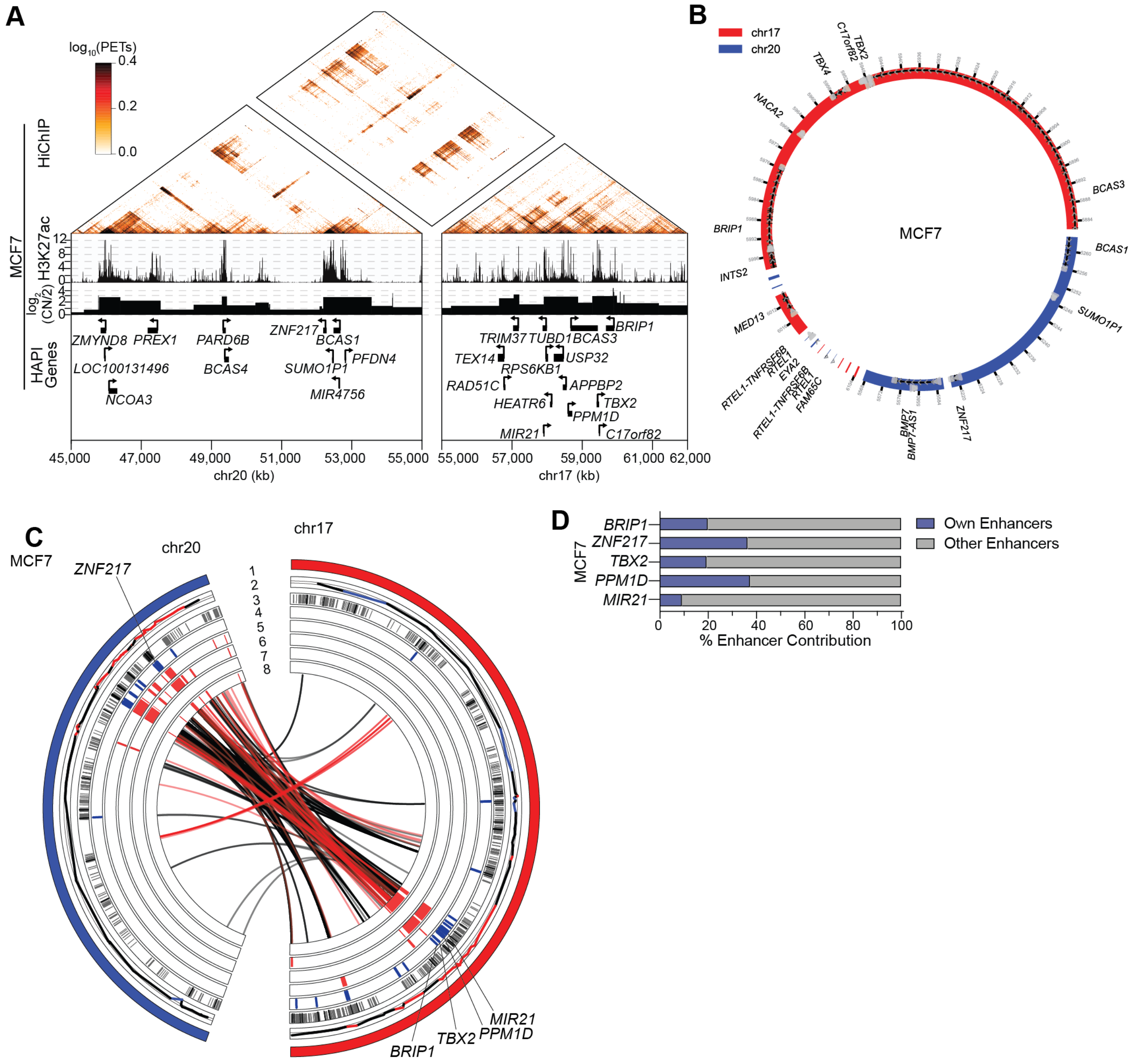

**Figure S7: A potential chimeric ecDNA event in MCF7. A:** H3K27ac HiChIP heatmap showing inter-chromosomal interactions between regions of chromosomes 17 and 20. **B:** Structure of a proposed ecDNA in MCF7 based on the AmpliconArchitect results. **C:** Circos plot of chromosomes 20 and 17 in MCF7 showing 1) chromosome, 2) copy number, 3) H3K27ac peaks showing enhancers (grey) and super-enhancers (black), 4) TSS of HAPI genes in MCF7, 5-8) DNA segments of AmpliconArchitect-predicted complex amplicons, and in the center HiChIP enhancer-promoter interactions of HAPI genes (black), and WGS translocations (red). **D:** Enhancer distribution of oncogenes harbored in the predicted complex amplicons in MCF7 cells. Enhancers within 2mb to a genes' promoter in the endogenous chromosomal locus will be considered as the gene's 'own' enhancers, otherwise will be considered as 'other' enhancers.

#### Supplementary Figure 8

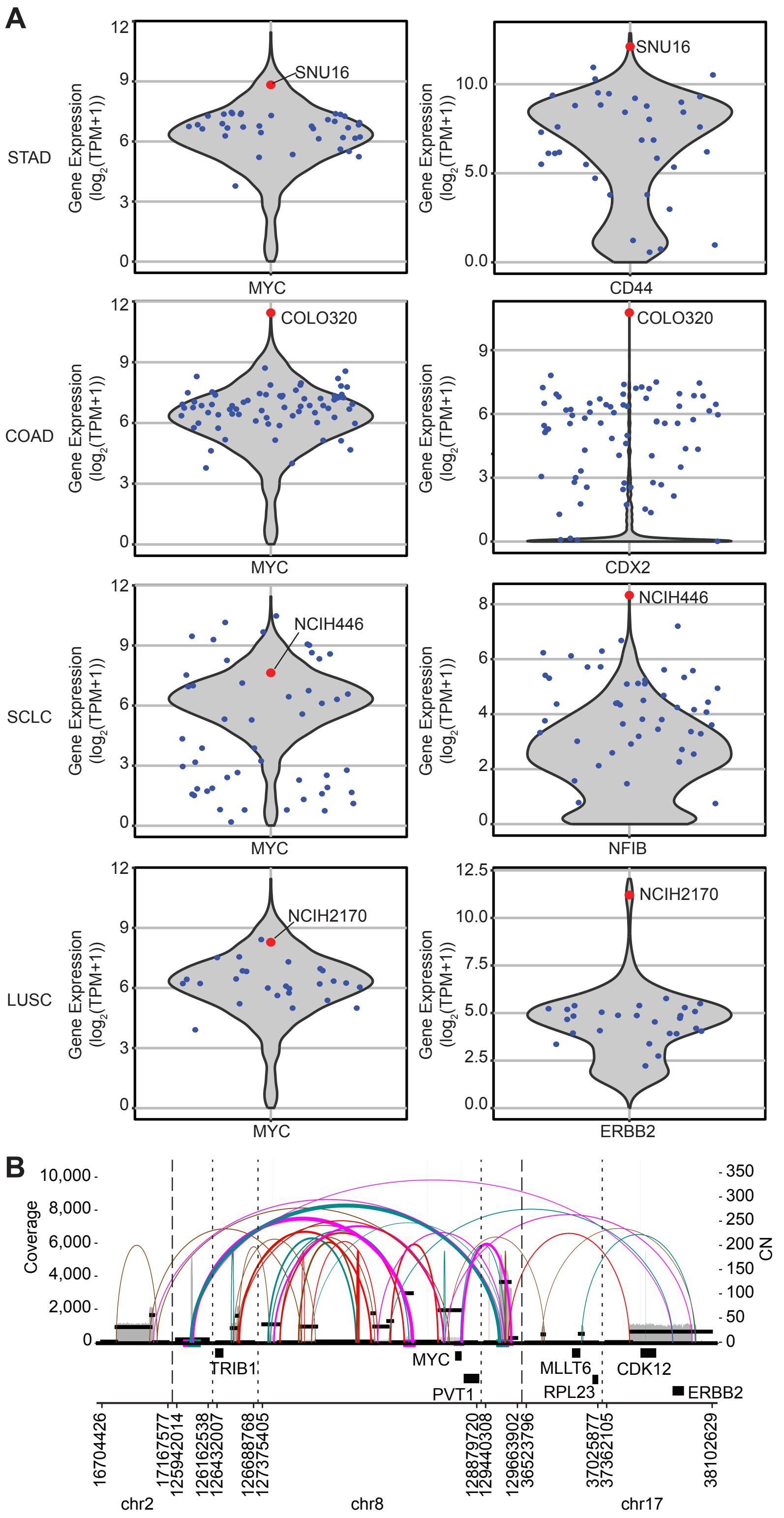


**Figure S8: Gene expression of ecDNA genes in cell lines of interest and profiling ecDNA in a PCAWG sample. A:** Expression of selected cancer-related genes present on ecDNAs in cell lines (red) representing STAD, COAD, SCLC, or LUSC compared to all other cell lines of the given cancer type (blue) and all cell lines in CCLE (grey violin plots). **B:** AmpliconArchitect reconstruction depicting WGS-derived copy number and structural variants for a PCAWG tumor sample containing the oncogenes *ERBB2* and *MYC*, similar to the structure found in NCIH2170.

#### Supplementary Figure 9

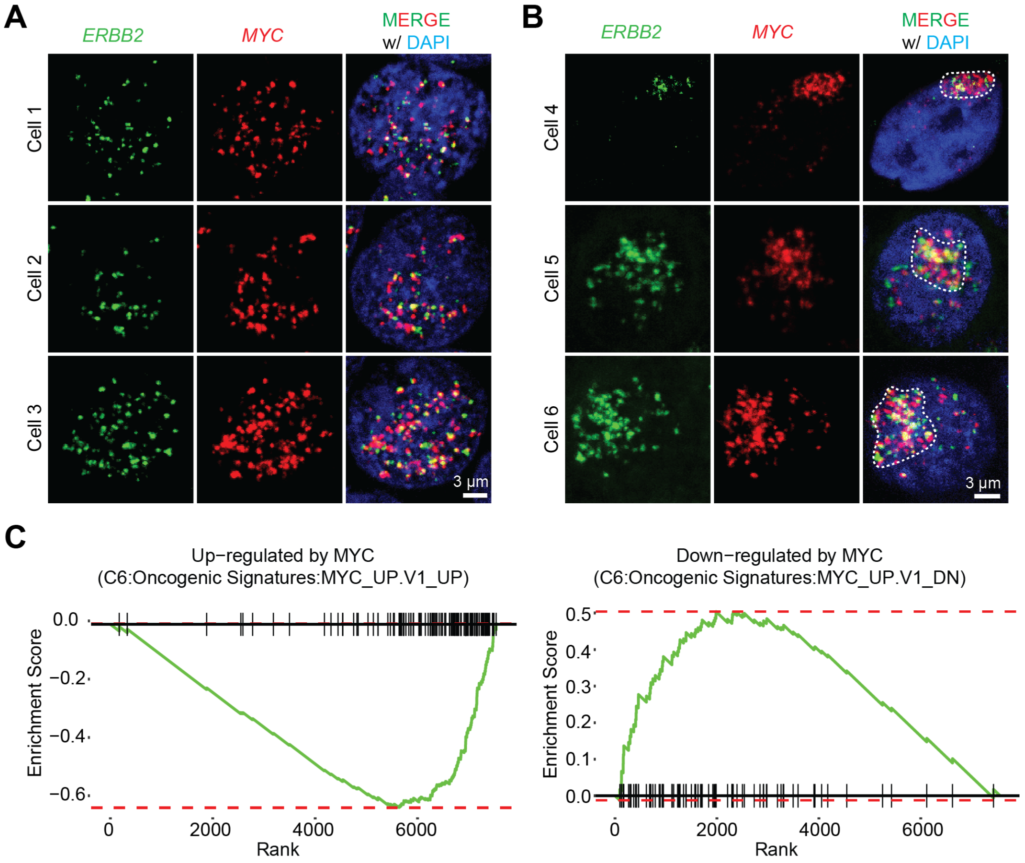


**Figure S9: Interphase DNA FISH imaging shows the spatial distribution of chimeric ecDNA. A:** Representative images of NCIH2170 cells in interphase showing the colocalization of *MYC* (red) and *ERBB2* (green) within cell nuclei (blue). In total, 219 interphase nuclei were collected from 8 separate images, which indicate the presence of high abundant *MYC-ERBB2* chimeric ecDNAs. **B:** The *MYC-ERBB2* ecDNAs can cluster into spatially segregated hubs in NCIH2170 cells. **C:** GSEA analysis measuring differential gene expression (CRISPRi *MYC* promoter vs non-targeting) of MYC up-regulated (left) and down-regulated (right) genes.

#### Supplementary Table 1: Cell lines and their associated omics data used in the study

| Cell Line | HiChIP Data | HiC-Pro allValidPairs | FASTQ File | Anchor Calling | H3K27ac ChIP-seq Data | WGS Data | AR ChIP-seq Data |
|---|---|---|---|---|---|---|---|
| LNCaP | GSE227242 | 37075641 | merged | broadpeak | GSE73994 | PRJNA361316 | GSE174109 |
| T47D | GSE227242 | 37390012 | single | broadpeak | GSE63109 | N/A | N/A |
| CUTLL1 | GSE228247 | 39244038 | merged | Self | N/A | N/A | N/A |
| REC1 | GSE228247 | 32423683 | merged | Self | GSE69558 | N/A | N/A |
| VCaP | GSE228247 | 20076305 | single | broadpeak | GSE105760 | N/A | N/A |
| MDAPCA2B | GSE228247 | 45838476 | merged | Self | N/A | N/A | N/A |
| 22RV1 | GSE228247 | 20175335 | single | broadpeak | GSE105627 | N/A | GSE94013 |
| ZR751 | GSE228247 | 51263859 | merged | broadpeak | GSE85158 | PRJNA523380 | N/A |
| MCF7 | GSE228247 | 59028122 | merged | broadpeak | GSE57436 | PRJNA523380 | N/A |
| MDAMB231 | GSE97585 | 37514192 | single | Self | N/A | N/A | N/A |
| MDAMB453 | GSE157381 | 37821066 | single | Self | N/A | N/A | N/A |
| DOHH2 | GSE228247 | 41769569 | merged | Self | GSE86743 | N/A | N/A |
| HT29 | GSE228247 | 26395716 | single | broadpeak | GSE73319 | N/A | N/A |
| HT55 | GSE147854 | 19478944 | single | Self | N/A | N/A | N/A |
| LK2 | GSE166232 | 13705192 | single | Self | N/A | N/A | N/A |
| H520 | GSE166232 | 20709304 | single | Self | N/A | N/A | N/A |
| DMS273 | GSE151002 | 36476957 | merged | Self | N/A | N/A | N/A |
| H69 | GSE151002 | 23465687 | single | Self | N/A | N/A | N/A |
| H146 | GSE151002 | 35660665 | merged | Self | N/A | N/A | N/A |
| H446 | GSE151002 | 39535685 | merged | Self | GSE115123 | PRJNA523380 | N/A |
| H2170 | GSE228247 | 37495342 | merged | Self | N/A | PRJNA523380 | N/A |
| SHP77 | GSE151002 | 21552164 | single | Self | N/A | N/A | N/A |
| COLO320DM | GSE159985 | 13880280 | single | Self | GSE159972 | PRJNA506071 | N/A |
| SNU16 | GSE159985 | 103392115 | single | Self | GSE159972 | PRJNA523380 | N/A |
| HMEC | GSE188401 | 35395988 | single | Self | N/A | N/A | N/A |
| PrEC | GSE188401 | 56483476 | single | Self | N/A | N/A | N/A |
| CORL88 | GSE151002 | 17946661 | single | Self | N/A | N/A | N/A |
| CORL279 | GSE151002 | 17010536 | single | Self | N/A | N/A | N/A |
| H524 | GSE151002 | 12298085 | single | Self | N/A | N/A | N/A |
| H1105 | GSE151002 | 18409434 | single | Self | N/A | N/A | N/A |
| H1694 | GSE151002 | 19689472 | single | Self | N/A | N/A | N/A |
| H1836 | GSE151002 | 19539922 | single | Self | N/A | N/A | N/A |
| H1876 | GSE228247 | 52514472 | merged | Self | N/A | N/A | N/A |
| H2029 | GSE151002 | 27050233 | single | Self | N/A | N/A | N/A |
| H2171 | GSE151002 | 18707959 | single | Self | N/A | N/A | N/A |
| H2066 | GSE151002 | 19674767 | single | Self | N/A | N/A | N/A |

**Supplementary Table 2: Pathway analysis for shared HAPI genes in each cancer type listed in Figure 1D**
(Note: only the top ranked pathway is listed in each category.)

| Cluster Groups | KEGG Legacy Pathways | | REACTOME | | Gene Ontology Molecular Function | |
|---|---|---|---|---|---|---|
| | **Pathway** | **FDR** | **Pathway** | **FDR** | **Pathway** | **FDR** |
| Blood | B-cell receptor signaling pathway | $4.74e^{-5}$ | Hemostasis | $2.53e^{-7}$ | Transcription regulatory activity | $6.81e^{-17}$ |
| BRCA | Systemic lupus erythematosus | $9.33e^{-13}$ | Signaling by Nuclear Receptors | $8.49e^{-21}$ | Transcription regulatory activity | $4.6e^{-20}$ |
| COAD | Pathways in cancer | $3.26e^{-11}$ | Developmental Biology | $2e^{-11}$ | Transcription regulatory activity | $4.39e^{-12}$ |
| LUSC | Pathways in cancer | $2.75e^{-6}$ | Developmental Biology | $5.73e^{-7}$ | Transcription regulatory activity | $4.01e^{-15}$ |
| PRAD | Systemic lupus erythematosus | $9.91e^{-12}$ | Signaling by Nuclear Receptors | $2.51e^{-20}$ | Transcription regulatory activity | $4.38e^{-26}$ |
| SCLC | Cell cycle | $2.85e^{-4}$ | Diseases of signal transduction by growth factor receptors and second messengers | $2.17e^{-6}$ | Transcription regulatory activity | $1.1e^{-12}$ |
| STAD | MAPK signaling pathway | $7.42e^{-4}$ | Nuclear Events (kinase and transcription factor activation) | $3.63e^{-3}$ | DNA binding transcription factor binding | $1.08e^{-10}$ |

#### Supplementary Table 3: Primers used in the study

##### sgRNAs for CRISPRi

| Name | Sequence | Blat Regions (hg19) | Notes |
|---|---|---|---|
| sg_NT1_F | CACCGCTGAGTGAAAAATAAAAGTT | N/A | Negative Control |
| sg_NT1_R | AAACAACTTTTATTTTTCACTCAGC | | |
| sg_NT2_F | CACCGATCGTTTCCGCTTAACGGCG | N/A | Negative Control |
| sg_NT2_R | AAACCGCCGTTAAGCGGAAACGATC | | |
| sg_ETV1_promoter_F | CACCGGGTCAGCAATAAACAACAA | chr7:14031070-14031090 | |
| sg_ETV1_promoter_R | AAACTTGTTGTTTATTGCTGACCC | | |
| sg_FOXA1_promoter_F | CACCGAGGGGACAATGAAGAGAAAC | chr14:38063490-38063509 | |
| sg_FOXA1_promoter_R | AAACGTTTCTCTTCATTGTCCCCTC | | |
| sg_LNCaP_e1_1_F | CACCGGAGAACTGGTCAAAAGGTA | chr14:37717096-37717115 | |
| sg_LNCaP_e1_1_R | AAACTACCTTTTGACCAGTTCTCC | | |
| sg_LNCaP_e1_2_F | CACCGAGGTATTAATACCAAGCCCC | chr14:37717126-37717145 | |
| sg_LNCaP_e1_2_R | AAACGGGGCTTGGTATTAATACCTC | | |
| sg_LNCaP_e2_1_F | CACCGCTTAATAAACTATTAAACTG | chr14:37798906-37798926 | |
| sg_LNCaP_e2_1_R | AAACCAGTTTAATAGTTTATTAAGC | | |
| sg_LNCaP_e2_2_F | CACCGTAAAACACAAAGCAGAGTTC | chr14:37798962-37798981 | |
| sg_LNCaP_e2_2_R | AAACGAACTCTGCTTTGTGTTTTAC | | |
| sg_LNCaP_e3_1_F | CACCGTTTCCTCCTCTATAAAAGTA | chr14:37887169-37887188 | |
| sg_LNCaP_e3_1_R | AAACTACTTTTATAGAGGAGGAAAC | | |
| sg_LNCaP_e3_2_F | CACCGCCAAGACCATGAATGGAAAT | chr14:37887324-37887343 | |
| sg_LNCaP_e3_2_R | AAACATTTCCATTCATGGTCTTGGC | | |
| sg_LNCaP_e4_1_F | CACCGCCCAGTCTTAAAAATACTGA | chr14:37905889-37905908 | |
| sg_LNCaP_e4_1_R | AAACTCAGTATTTTTAAGACTGGGC | | |
| sg_LNCaP_e4_2_F | CACCGAAAATAACAATGTTAAGTCC | chr14:37905937-37905956 | |
| sg_LNCaP_e4_2_R | AAACGGACTTAACATTGTTATTTTC | | |
| sg_CCND1_promoter_F | CACCGCGAGGGGCAGAAGAGCGCGA | chr11:69455974-69455993 | |
| sg_CCND1_promoter_R | AAACTCGCGCTCTTCTGCCCCTCGC | | |
| sg_ZR75_e1_F | CACCGCTCATGGAGAAAAGGAACAA | chr8:29380258-29380277 | |
| sg_ZR75_e1_R | AAACTTGTTCCTTTTCTCCATGAGC | | |
| sg_ZR75_e2_F | CACCGTAACAGCTTAAAGGTTATTG | chr8:29387600-29387619 | |
| sg_ZR75_e2_R | AAACCAATAACCTTTAAGCTGTTAC | | |
| sg_ZR75_e3_F | CACCGACCTGGTAAAGTTCAACTGA | chr8:29596128-29596147 | |
| sg_ZR75_e3_R | AAACTCAGTTGAACTTTACCAGGTC | | |
| sg_ZR75_e4_F | CACCGCAGGAATTTCTTATTTGCAT | chr8:29614509-29614528 | |
| sg_ZR75_e4_R | AAACATGCAAATAAGAAATTCCTGC | | |
| sg_ZR75_e5_F | CACCGGACAGGTGACCCTGCCTGC | chr8:29631787-29631806 | |
| sg_ZR75_e5_R | AAACGCAGGCAGGGTCACCTGTCC | | |
| sg_MYC_promoter_sg1_F | CACCGTAATTCCAGCGAGAGGCAG | chr8:128748488-128748507 | |
| sg_MYC_promoter_sg1_R | AAACCTGCCTCTCGCTGGAATTAC | | |
| sg_MYC_promoter_sg2_F | CACCGCAGCGCAGCTCTGCTCGCC | chr8:128748546-128,748565 | |
| sg_MYC_promoter_sg2_R | AAACGGCGAGCAGAGCTGCGCTGC | | |

##### RT-qPCR Primers

| Name | Sequence | Target Gene | Notes |
|---|---|---|---|
| RT_CTCF_F | CCCACACCGGGGAGAAGCCT | CTCF | Loading control |
| RT_CTCF_R | CGCCATCTGGGCCAGCACAA | | |
| RT_ETV1_F | CTTAGCCGTTCACTCCGCTA | ETV1 | |
| RT_ETV1_R | TCCTCCTCGTTGATGTGACG | | |
| RT_CCND1_F | TACTACCGCCTCACACGCTT | CCND1 | |
| RT_CCND1_R | CTTGGGGTCCATGTTCTGCTG | | |
| RT_DUSP4_F | GGCATCACGGCTCTGTTGAA | DUSP4 | |
| RT_DUSP4_R | TGTCGGCCTTGTGGTTATCT | | |
| RT_FOXA1_F | CTACTACGCAGACACGCAGG | FOXA1 | |
| RT_FOXA1_R | CCGCTCGTAGTCATGGTGTT | | |
| RT_TTC6_F | CACTGATAAGCCGGACTAACG | TTC6 | |
| RT_TTC6_R | ATTTCCCCGTCCAACATAAGC | | |
